# Spectral patterns of MEG oscillatory coupling emerge from meta-stable dynamics with small coupling delays

**DOI:** 10.1101/2025.08.18.670813

**Authors:** Felix Siebenhühner, Francesca Castaldo, Joana Cabral, J. Matias Palva, Gustavo Deco, Satu Palva

## Abstract

Functional connectivity (FC) is a fundamental mechanism of neural communication, connecting distinct oscillating populations. Oscillatory networks exhibit heterogeneity across frequencies and coupling modes whose origins are not well understood, but have been suggested to involve a complex interplay of critical-like dynamics and structure-function coupling. We here utilized structural connectivity (SC) to tune a whole-brain computational model of delay-coupled damped oscillators near a Hopf bifurcation to match oscillations and FC as observed in resting-state magnetoencephalography (MEG) data.

We assessed two forms of oscillation-based FC from empirical and model data, namely phase synchronization (PS) and amplitude coupling (AC). We found that both oscillations and FC best matched with empirical observations in a meta-stable regime which was characterized by small delays, realistic oscillation lifetimes, and FC with intermediate strength and high variability. How well MEG FC patterns were matched by the model varied between frequency bands and best fits were observed for high-alpha and beta-band networks. These fits could partially, but not fully, be explained by correlations with SC, implicating that both structure-function coupling and critical-like metastable dynamics underlie empirical FC, and the contributions of these mechanisms varies between different frequency bands in MEG data.

## Introduction

Brain activity measured with functional MRI (fMRI) is organized into coherent functional networks both during task and rest (Deco et al., 2011; Dosenbach et al., 2007; Fox et al., 2005; Raichle, 2015) which show spatio-temporal variability on different time scales. On a coarse time scale, these networks have been assessed with functional magnetic resonance imaging (fMRI), from which functional connectivity (FC) and its dynamic - time varying - patterns can be derived (Griffa et al., 2017; Vidaurre et al., 2017). It has been established that BOLD FC network dynamics are constrained by the brain’s structural connectome (Griffa et al., 2022; Preti & Van De Ville, 2019; Sadaghiani & Wirsich, 2020) and microstructure (Basile et al., 2022; Larivière et al., 2019; McDonough & Siegel, 2018; Teipel et al., 2010) in a region-specific manner. This structure-function coupling (SFC) has been found to vary between individuals and age (Baum et al., 2020) and decrease from unimodal regions low in the processing hierarchy towards the higher association regions (Liu et al., 2023).

However, at the neurophysiological mechanistic level, measured with electrophysiological methods, collective neuronal activity is organized by rhythmic excitability fluctuations — transient or dampened neuronal oscillations — that are abundant across the brain’s processing hierarchy and spatio-temporal scales. Oscillations and their inter-areal coupling are thought to provide a temporal clocking mechanism for dynamic routing of neuronal information processing via oscillatory ensembles codes (Engel et al., 2001; Fries, 2015; Hahn et al., 2019; Siegel et al., 2012; Singer, 1999; Vinck et al., 2023). In humans, oscillations can be measured non-invasively with electro- and magnetoencephalography (EEG/MEG). Individual source reconstruction of M/EEG data has been used to establish that oscillation dynamics and their large-scale spatio-temporal patterns across oscillatory frequencies are fundamental for various behaviors and higher cognitive functions (Engel et al., 2001; Fries, 2015; S. Palva & Palva, 2012; Sauseng & Klimesch, 2008; Siegel et al., 2012; Singer, 1999; Womelsdorf et al., 2007). These oscillatory dynamics exhibit, however, large heterogeneity which is influenced by genetic factors (Simola et al., 2022), neurotransmitter microstructure (Siebenhühner et al., 2024), and individual variability in brain states that has been linked to brain criticality (Fuscà et al., 2023). In contrast to fMRI, few studies have investigated how M/EEG FC is influenced by structure. While there are multiple models explaining the emergence of oscillatory coupling at micro-circuit level (Jones et al., 2007; Traub et al., 2004; Whittington et al., 1995), it is not fully understood how large-scale brain networks emerge from these micro-scale dynamics.

In order to understand causal network dynamics at the macroscopic level, whole-brain computational models constrained by empirical structural connectivity data can be leveraged. A decade of pioneering work has established that fMRI-derived large-scale brain networks can be approximated with various whole brain computational models, such as Wilson-Cowan oscillators (Deco et al., 2009; Traub et al., 2004; Whittington et al., 1995), mean-field (Deco et al., 2014), Kuramoto (Cabral et al., 2011), and Hopf bifurcation models (Deco, Kringelbach, et al., 2017). Several of these studies have provided evidence that brain network dynamics exhibit metastable, and thus critical-like, dynamics (Deco, Kringelbach, et al., 2017; Deco & Jirsa, 2012; E. C. A. Hansen et al., 2015) which is also supported by studies of fMRI data by itself (beim Graben et al., 2019; Hancock et al., 2022). These findings complement other pioneering experimental (Fuscà et al., 2023; Kinouchi & Copelli, 2006; Tagliazucchi et al., 2012) and theoretical work (Chialvo, 2010; Cocchi et al., 2017) in establishing that the healthy brain operates near the critical transition at balanced excitation / inhibition (E/I) between subcritical (dampened) and supercritical (hyper-synchronized) phases in the system’s state space characterized by large variance, scale-freeness, and long-range spatiotemporal correlations.

Contrasted with fMRI FC, there have been fewer studies leveraging whole-brain computational models to understand M/EEG-derived FC (Cabral et al., 2014, Cabral et al., 2017; Deco, Cabral, et al., 2017). A recent advance has been the introduction of a whole-brain model of coupled oscillators operating near a Hopf bifurcation which incorporates both empirical SC as connections strengths and delays in large-scale network dynamics to model oscillations in the presence of noise (Cabral et al., 2022; Castaldo et al., 2023; Ponce-Alvarez & Deco, 2024). This model uses coupled Stuart-Landau oscillators operating in the damped subcritical regime of a Hopf bifurcation. Importantly, incorporating non-zero and variable coupling delays informed by SC produces meta-stable modes of dampened (or transient) oscillations (Cabral et al., 2022; Castaldo et al., 2023). These dynamics match empirical observations of cortical oscillations being mostly short lived, with their lifetimes varying by region and frequency band (Freyer et al., 2009; Myrov et al., 2024; Quinn et al., 2019; van Ede et al., 2018).

We here used this model to provide mechanistic understanding of whether empirical MEG oscillation-based FC emerges in meta-stable critical brain modes and how it is influenced by time delays arising from SC. We first assessed the presence of transient oscillations via estimation of their lifetimes with the novel phase autocorrelation function (Myrov et al., 2024) for both MEG and model data to constrain the model to realistic coupling strengths and inter-areal delays. As FC patterns in MEG data have been shown to differ between coupling modes (Engel et al., 2013; Siebenhühner et al., 2024; Siems & Siegel, 2020), we here assessed two modes, inter-areal phase synchrony (PS) and amplitude correlations (AC), for both MEG and model data. Assessing the correspondence of spatial FC patterns between MEG and model, we found that the model produced the best fits for empirical oscillatory FC in high-alpha (13–15 Hz) and beta (15–30 Hz) bands, where model FC emerges in meta-stable coupling models in a meta-stable regime with high variability. MEG FC could only be partially explained by SC, implying that critical dynamics contribute significantly to the spatial architecture of FC. Our study thus provides evidence that oscillatory coupling as observed in MEG emerges from the brain operating at near-critical meta-stable dynamics and the interplay of critical-like dynamics with structural constraints in a frequency-specific manner.

## Results

### Transient oscillations and functional connectivity in MEG data and Hopf model

We first assessed oscillation-based FC from individually source-reconstructed resting-state MEG data from N = 60 healthy human subjects. The signals in 100 cortical parcels of the Schaefer atlas were filtered at 41 different, log-spaced, narrow frequency bands, *f*_*filt*_ from 1 to 96 Hz (Figure 1A-C). Group-mean phase-autocorrelation (pACF) lifetimes, which have been recently introduced to assess the lifetimes of transient oscillations (Myrov et al., 2024), exhibited a main narrow peak in the alpha-band at around *f*_*filt*_ = 10 Hz showing transient rhythmicity for 3.5 cycles on average, and weaker rhythmicity in beta (15 - 30 Hz) and gamma (30 - 100 Hz) bands (Figure 1B). We estimated functional connectivity (FC) between all parcel pairs for two coupling modes - phase synchrony (PS) and amplitude coupling (AC), which were estimated with the imaginary phase-locking value (iPLV) and orthogonalized correlation coefficient (oCC), respectively, that are insensitive to direct effects of signal leakage in MEG source data (J. M. Palva et al., 2018). The group mean PS and AC showed robust FC in theta-beta range (4 - 30 Hz) peaking at *f*_*filt*_ = 10 Hz (Figures 1C, 2A-B), as reported previously (Fuscà et al., 2023; Siebenhühner et al., 2024).

**Figure 1.**
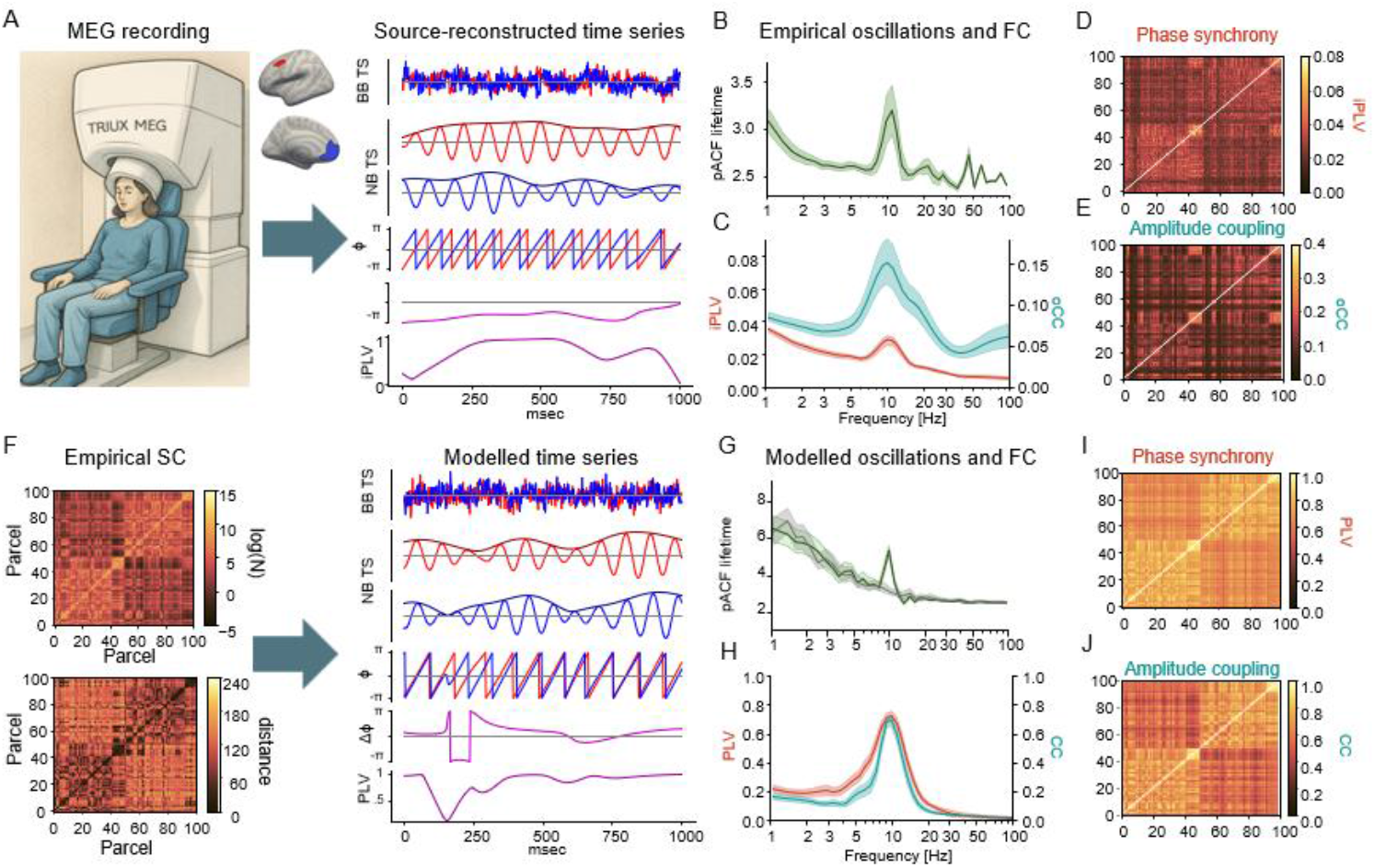
Transient oscillations and functional connectivity in MEG and model data. **A**. Resting-state MEG data is source-reconstructed and the resulting broadband (BB) data is filtered using Morlet-wavelets to obtain narrowband (NB) data in 41 frequencies. Example of broad band (top row), and alpha band (11Hz) filtered traces (2nd, 3rd row) for parcels of left and right prefrontal cortex (red: DM_PFC_6_L, blue: DM_PFC_3_L) and their phase time series (TS) (4th row), phase difference, and imaginary Phase Locking Value (iPLV). **B**. Group-mean (N=60) oscillation lifetimes assessed with pACF as a function of filtering frequency. **C**. Grand average phase synchrony (estimated with iPLV) and amplitude coupling (est. with oCC) averaged over subjects and edges. **D–E**. Group-mean pairwise phase synchrony and amplitude coupling between all 100 parcels at 11 Hz. **F**. Using empirical coupling strengths and delays as input, broadband time series are simulated for Stuart-Landau oscillators (this and following panels with parameters: global coupling strength K=15.9, mean delay MD = 5 msec, bifurcation parameter a = −5) and then filtered as done for MEG data. **G**. Mean pACF lifetimes. Green: modelled data, grey: surrogates. **H**. Mean phase synchrony and amplitude coupling. **I, J**. Pairwise phase synchrony and amplitude coupling between all 100 oscillators at 11 Hz.

We used a whole-brain model of coupled Stuart-Landau oscillators near a Hopf bifurcation to simulate brain activity for 50 seconds in 100 nodes corresponding to the 100 brain parcels of the Schaefer atlas. Human tractography data from the human connectome project <u>(HCP</u>) was used to determine coupling strengths and delays (from mean fiber numbers and strengths, resp.) between nodes in the model (Supp. Figure 1). We ran simulations of the model for multiple combinations of coupling strength *K*, mean delay *MD*, natural oscillator frequency *f*_*nat*_ and bifurcation parameter *a*. In each run, all 100 oscillators had the same values for *f*_*nat*_ and *a*, so that regional specificity was determined by connectivity alone. We then assessed oscillation lifetimes in the modelled data with pACF as well as PS and AC, but using the phase-locking value (PLV) and correlation coefficient (CC) as there is no signal leakage in the model data (Figures 1D-F). The filtering frequencies *f*_*filt*_ at which peak lifetimes were observed shifted as a function of K, MD, and *f*_*nat*_ – whereas varying the bifurcation parameter a had little influence on lifetimes except at MD = 1 msec (Figure 1G, Supp. Figures 2-3). Notably, even at low coupling strengths, oscillation lifetimes in the coupled model always peaked at frequencies below the oscillators’ natural frequencies. Realistic lifetimes that were similar to those found for oscillations in MEG were observed at mean delays (MD) of 3 or 5 msec and moderate coupling strengths (Figure 2G, Supp. Figure 2). Therefore we tuned the model to operate with these parameter values to understand the emergence of large-scale oscillatory dynamics.

**Figure 2.**
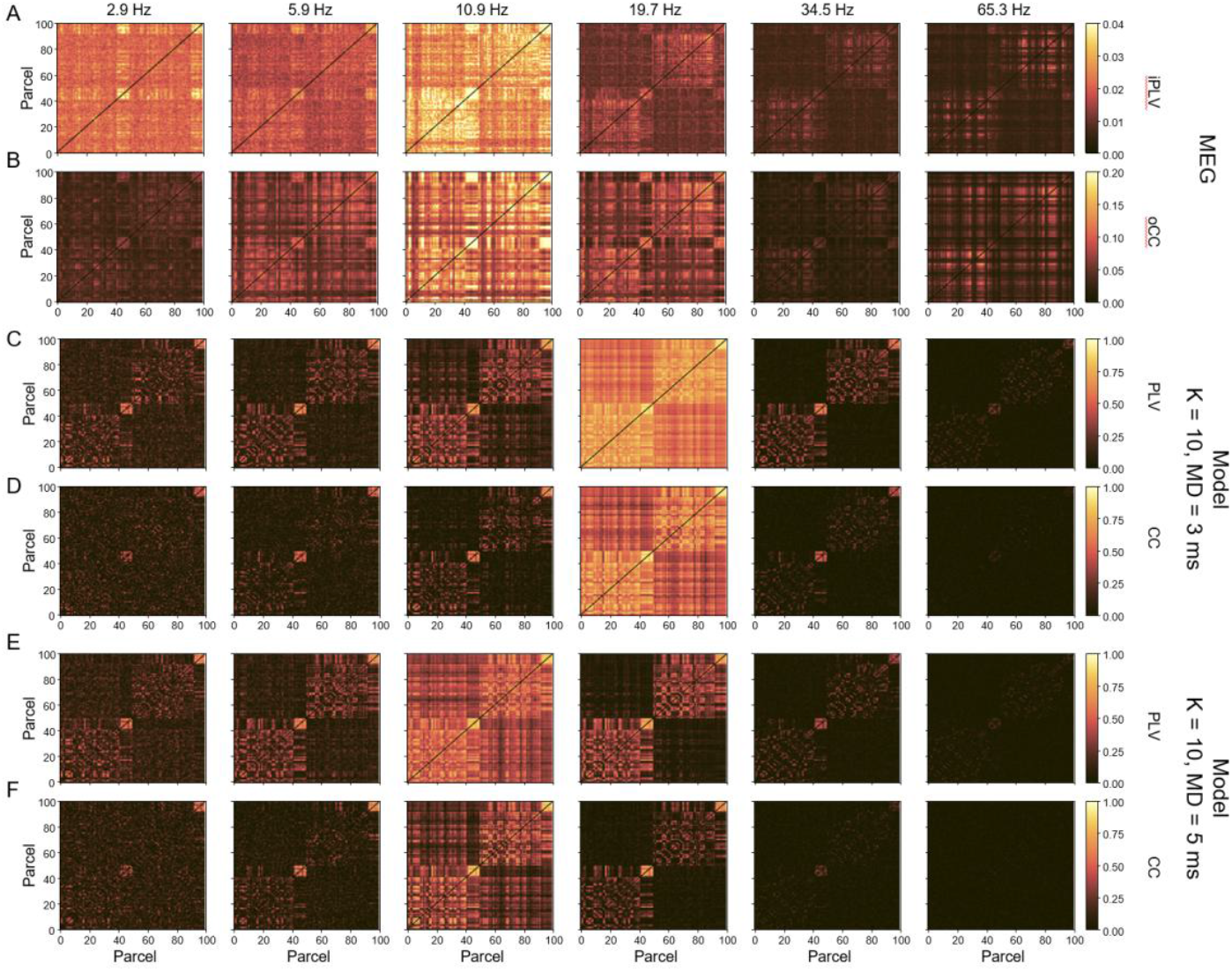
FC varies with frequency and with model parameters. **A**,**B**. MEG-derived FC between all 100 parcels averaged across participants for A. mean phase synchrony (PS) estimated with iPLV, and B. amplitude correlations (AC), estimated with OCC, at selected filtering frequencies. **C**,**D**. Model FC at K = 10, MD = 3 msec, f_nat_ = 40 Hz and a = −5 for C. PS (PLV) and D. AC (CC). **E**,**F**. Same as in C-D but for K = 10 and MD = 5 msec

### The Hopf model exhibits a critical transition from hypo-to hypersynchronization

We then assessed variability in the PS and AC at the edge level for representative frequencies. In the MEG data, PS and AC values (assessed with iPLV and oCC, respectively) were moderate and showed variability across anatomically confined systems (Figure 2A-B). In the model, spatial patterns of FC resembled those of MEG, but coupling strength (PS and AC assessed with PLV and CC, resp.) varied more with filtering frequency and approached values of 1 at specific frequencies, depending on model parameters. For realistic mean delay MD of 3 or 5 msec (Figure 2C-F) and medium coupling strength K = 10, this “hypersynchronization” was observed for beta and alpha band, respectively. In the parameter space spanned by all possible values of MD, K, this hypersynchronization was observed in a contiguous region spanning from filtering frequencies in low gamma band (up to the natural frequency of 40 Hz of the model’s Stuart-Landau oscillators) at low K and MD to delta band for very high K and MD. As had been the case for oscillation lifetimes (Supp Fig S2A), no synchronization was observed in frequencies above 40 Hz (Figure 3 A,C). Generally, hypersynchronization zones appeared somewhat wider for PS then AC.

**Figure 3.**
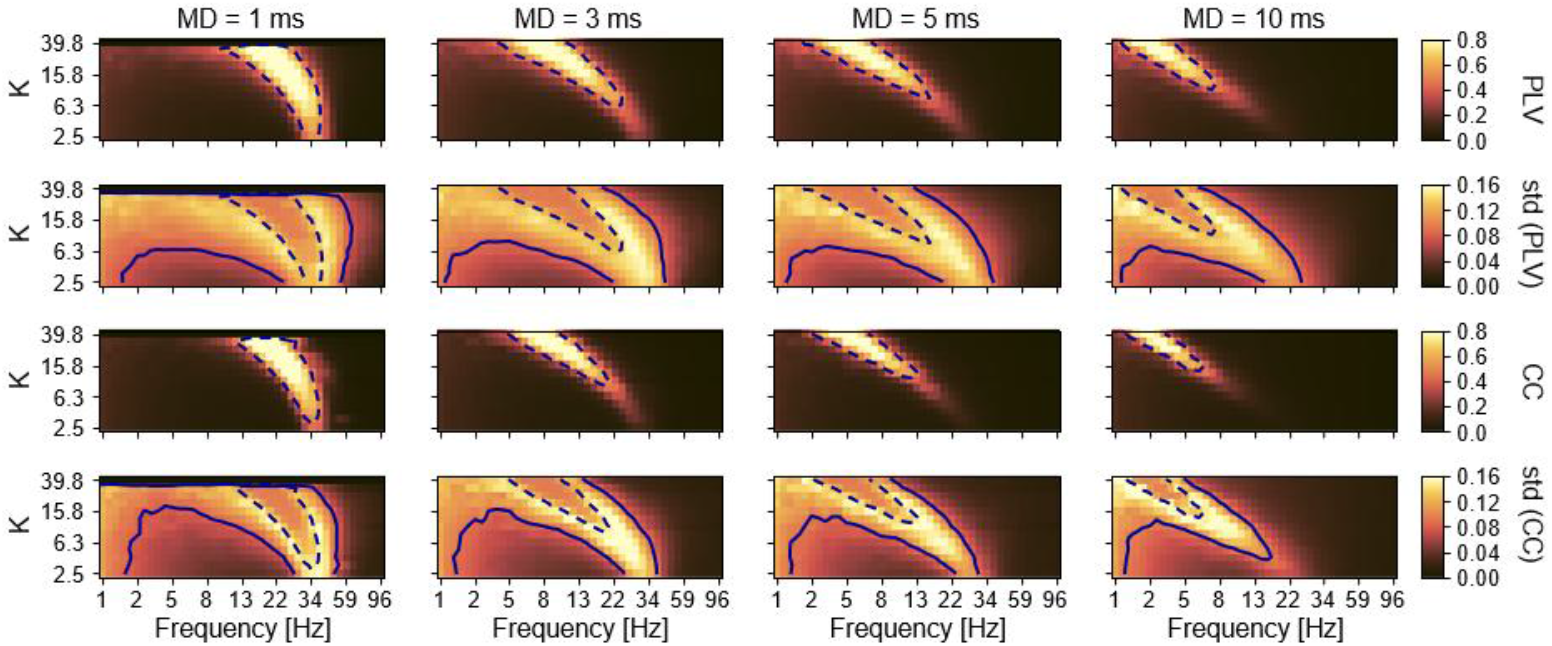
Metastable dynamics occur at moderate FC. **A**. Mean and **B**. standard deviation of phase synchrony (PLV) over all edges as a function of filtering frequency (x-axis), coupling strength K (y-axis), mean delay MD (column), for f_nat_ = 40 Hz and a = −5. Dashed contours indicate the median value (across all values on the heatmap) of the mean, the solid contours in B indicate the median value of the standard deviation. **C**. and **D**. Same for amplitude correlations (CC).

### Model FC exhibit critical metastable dynamics

To then assess the presence of metastable critical-like dynamics in FC, we assessed the variability in PS and AC with the standard deviation (SD). High variability, indicating the presence of meta-stability, was observed in regions of intermediate coupling levels in the vicinity of the hyper-synchronized regions, and thus in frequencies just below and above those where synchronization was highest for any given combination of K and MD parameter values. The frequencies with the highest variability also depended on model parameters. For realistic delays of 3 or 5 msec, maximum variability occur for very high K, but was strongest in frequencies above the hypersynchrony frequencies, i.e. in the beta-band (Figure 3B,D, Supp. Figure 4).

In order to compare the range of hypersynchronized and metastable dynamics between PS and AC which are not comparable in absolute values, we then computed time-shuffled surrogate data for both in order to estimate the fraction of connections that were significant (*p* < 0.01) against surrogate values. In the hyper-synchronized zone, this fraction was 1 for both PS and AC, however both this and the metastable zone were wider in in PS (Supp. Figure 5).

We then set out to investigate the influence of the other modelling parameters. As with pACF lifetimes, FC was little affected by varying the bifurcation parameter *a*, except at MD = 1 and high *a*, where hypersynchronization was observed much more commonly (Supp. Figure 6). However, and again similar to pACF lifetimes, regions of high FC scaled with the oscillators’ natural frequency (Supp. Figure 7).

### MEG-derived FC is predicted by metastable critical dynamics in the model

To assess if empirical FC in source-reconstructed MEG data can be predicted by model FC specifically at the metastable critical dynamics, we estimated fits between model FC and MEG FC by computing the correlation of FC values across parcel pairs. The best fits were observed for f_filt_ in the high-alpha (13 – 15), beta (15 - 30 Hz), and low-gamma (30 – 40 Hz) bands in the high-variability regions of parameter space, and i.e. thus where metastability was highest (Figure 4A).

**Figure 4.**
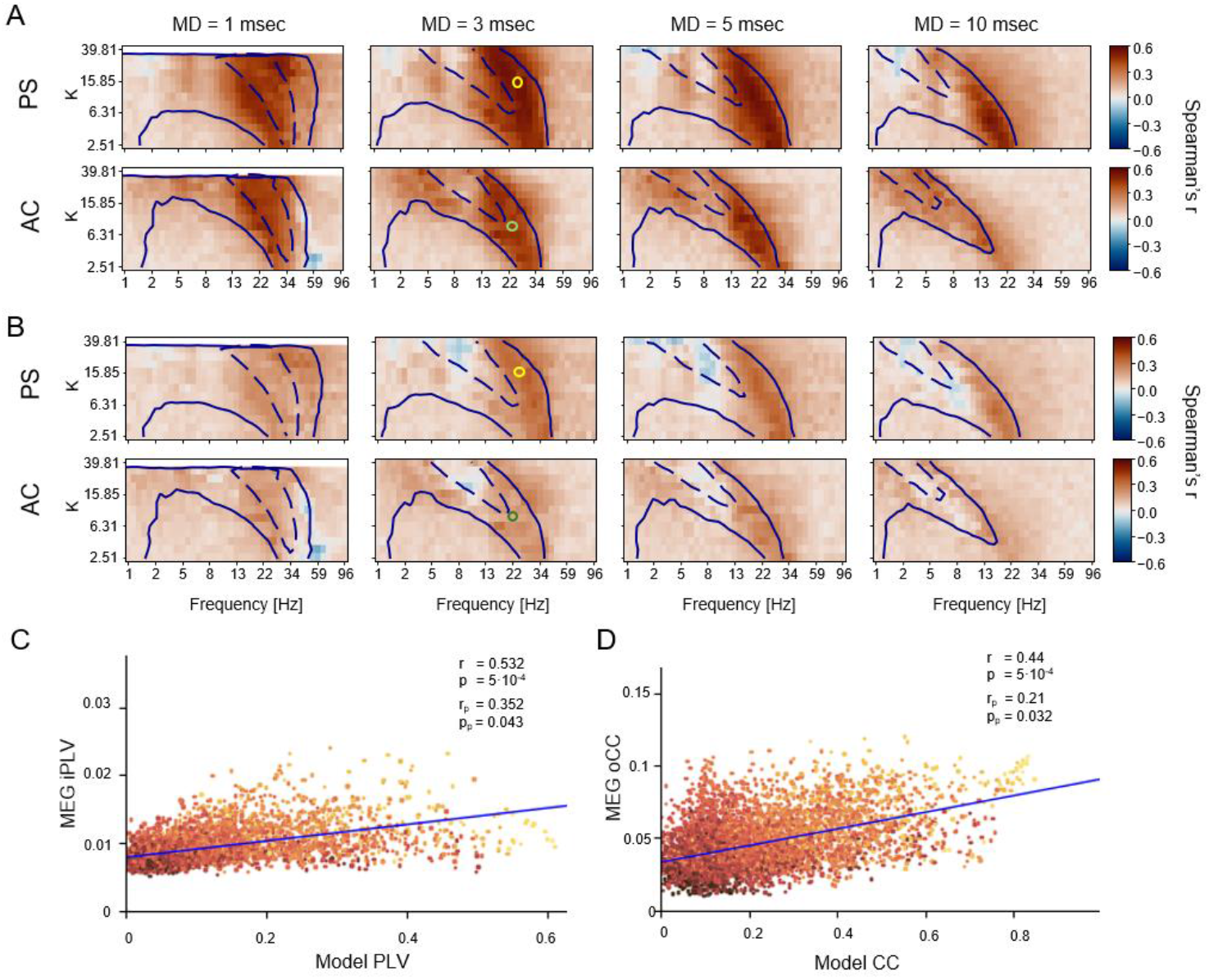
MEG functional connectivity is predicted by model FC in high-alpha, beta bands. **A**. Fits (Spearman’s r) between model FC and group-mean MEG FC for different values of coupling strength K, mean delay MD, and filtering frequency. Dashed and solid contours indicate the median values of the mean and standard deviation, resp., of model FC, as in Figure 3. **B**. Partial correlation between model FC and group-mean MEG FC, with SC as a covariate factor, same contours as in A. Right: Same for model AC and MEG AC. **C**. Scatterplot of modelled and MEG PS; each dot representing one edge. Model parameters: K = 15.8, MD = 3 msec, f_filt_ = 28.7 Hz (as indicated in A, B by the yellow circles); with r indicating the Spearman’s correlation, and p the p-value from comparison against spin permutations (N_perm_ = 2,000); r_p_ indicating the partial Spearman’s correlation and p_p_ the corresponding p-value (against spin permutations). Dot color represents the logarithm of the structural connectivity strength (log-value range −4 to 11). **D**. Scatterplot of modelled and MEG AC at K = 10, MD = 3 msec, f_filt_ = 23.7 Hz, corresponding to the green circles in A, B.

To estimate how the underlying SC contributed to MEG-FC prediction, we computed these fits using SC as a covariate. Correlations between model FC and MEG FC were reduced, but still significant for beta band albeit not for alpha (Figure 4B), indicating that MEG FC cannot be explained solely directly from SC, but that emergent critical metastable dynamics also contribute. In contrast, strong correlations between modeled FC and SC were observed at both hypersynchronized regions and high-variability regions (Supp Figure 8).

To more precisely understand how the model explained experimental MEG FC dynamics, we evaluated the fits between model and MEG FC matrices across filtering frequencies. Unlike in the preceding analysis, this meant that we now correlated model and MEG FC matrices that had been estimated between parcel time series filtered at different Morlet frequencies (i.e., we correlated MEG FC at filtering frequencies f_filt (MEG)_ with model FC at filtering frequencies f_filt (model)_). The quality of fits between model FC and MEG FC depended mostly on the *MD* and the filtering frequency of model time series, but less on the frequency at which MEG time series were filtered. The best fit values were consistently observed with MEG alpha-beta-band FC regardless of the model frequency indicating that FC in this range can be best explained by metastable dynamics with high variability in this model. In comparison, in addition to beta-band, there were also moderate correlations with MEG delta-theta (1-9 Hz) band FC (Figure 5) that persisted, albeit with reduced correlation coefficients, after controlling for SC as a covariate (Supp. Figure 9). These findings provide further support for the view that while MEG-derived PS and AC share similar characteristics, they are due to partially distinct mechanisms, in line with previous findings (Engel et al., 2013; Siebenhühner et al., 2024; Siems & Siegel, 2020).

**Figure 5.**
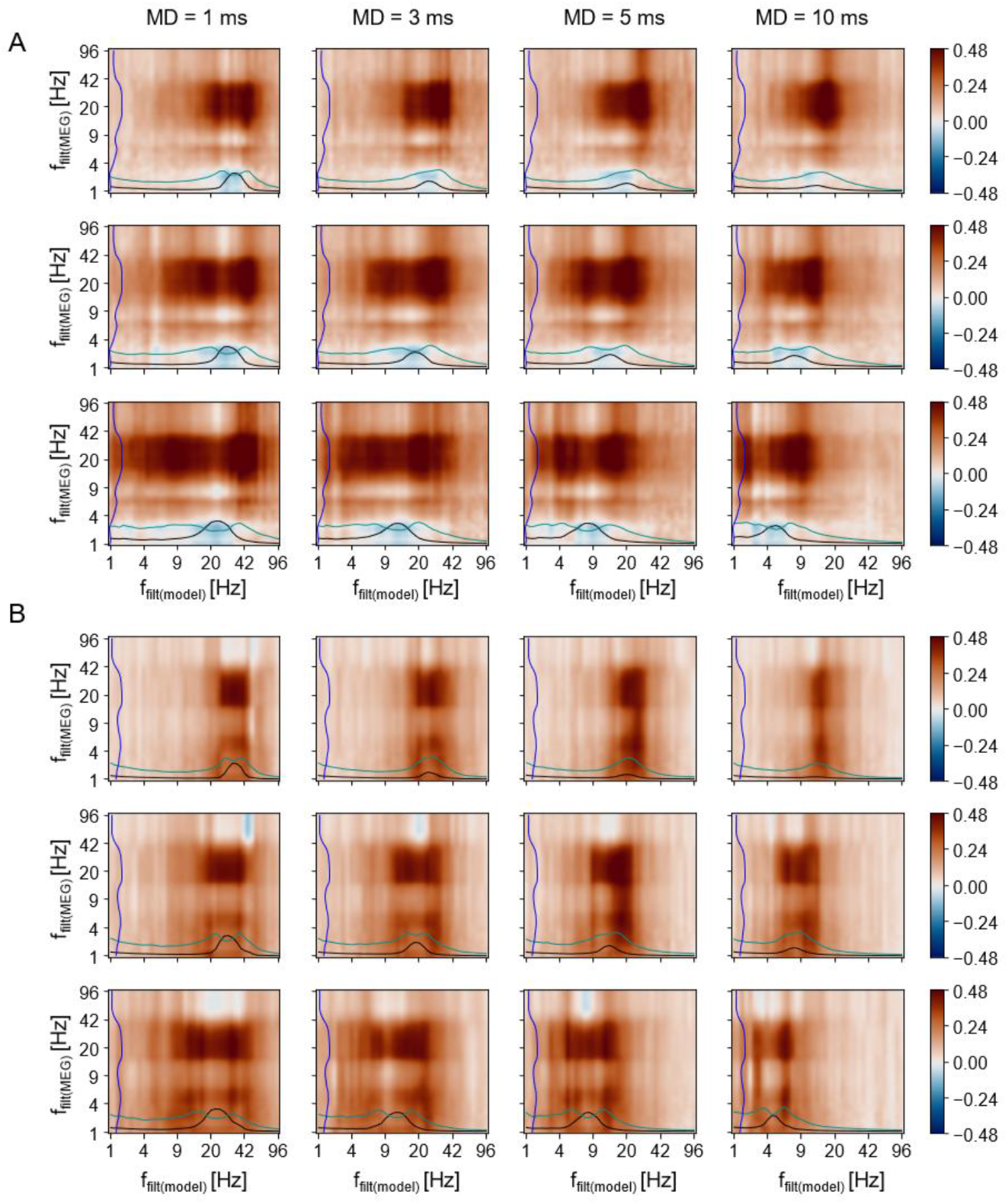
MEG beta-band connectivity correlates strongest with model connectivity across filtering frequencies. **A**. Correlations (Spearman’s r, across all edges) between model PS and group-mean MEG PS when both are filtered at different Morlet frequencies (x-axis: model and y-axis: MEG) for different values of coupling strength K (rows) and mean delay MD (columns). Black and cyan lines: mean and standard deviation of PS in the model as a function of frequency (a.u.); blue line: correlation of MEG PS with SC (a.u.). **B**. Same for AC.

## Discussion

Computational models have contributed to our understanding of brain function and the mechanisms underlying emergence of functional connectivity (FC) by assessing the influence of structural connectivity (SC) and model parameters that correspond to characteristics of the human brain (Deco et al., 2013; Ponce-Alvarez & Deco, 2024). Recent modelling work reproducing FC patterns from neuroimaging data has provided evidence that FC exhibits characteristics indicative of critical dynamics such as metastability (Cabral et al., 2022; Castaldo et al., 2023; Deco, Kringelbach, et al., 2017; Deco & Jirsa, 2012; E. C. A. Hansen et al., 2015; Tognoli & Kelso, 2014) and that it is partially constrained by structural connectivity (SC) via structure-function coupling (SFC) (). We here used the whole-brain Hopf model of delay-coupled Stuart-Landau oscillators (Cabral et al., 2022) to simulate realistic neuronal time series and evaluated the fit of oscillations and FC in the model to source-reconstructed MEG data. Our results show that damped oscillations and metastable FC result from incorporating coupling strengths and realistic delays informed by SC, in line with previous findings (Baum et al., 2020; Castaldo et al., 2023; Liu et al., 2023).

It has recently been established in theoretical frameworks and computational models (Lundqvist et al., 2024; Torres et al., 2024) that local oscillations in the brain tend to be transient or ‘bursty’ rather than continuous (Tal et al., 2020). The majority of evidence so far arise from local field potential (LFP) data in animal models (Buzśaki & Wang, 2012; Lundqvist et al., 2016) while evidence is scarce for EEG/MEG, where oscillations typically decay over a ‘lifetime’ in the range of 3 - 8 cycles rather than being sustained (Myrov et al., 2024; Stokes et al., 2023). We here modelled oscillations with the Hopf model in which Stuart-Landau oscillators below the bifurcation limit in the subcritical regime show damped, or slowly decaying, oscillations when perturbed by noise and/or input from coupled oscillators, in contrast to the supercritical case where they exhibit limit-cycle oscillations (Cabral et al., 2022). Further, the model performs better when using realistic, individual delays between regions informed by SC rather than homogenous delays (Castaldo et al., 2023). Here we tuned the model to match empirical findings of burstiness of oscillations. We then showed causal evidence from the model that MEG functional connectivity arises from transient oscillations produced in a model by intermediate structural connectivity delays. In contrast, in networks with smaller delays, lifetimes were unrealistically long, and with larger delays, excessively short. This is in line with earlier findings from computational models that coupling delays shape the frequency spectrum of coupled oscillatory systems and higher delays and correlate inversely with oscillation amplitudes (Cabral et al., 2022; Castaldo et al., 2023). Further, stronger coupling not only increased estimated FC, but also slowed the oscillators’ effective frequency below their natural frequency, again confirming previous findings that delay-coupled oscillations exert a strong influence on each other via synchronization and coupling mechanisms.

### Critical metastability observed in both phase and amplitude coupling

Previous research has shown that large-scale brain activity shows signs of operating near a critical phase transition (Chialvo, 2010; Cocchi et al., 2017; Haldeman & Beggs, 2005). Critical dynamics are thus a macro-level emergent property of the brain that confer advantages such as maximal dynamic range (Kinouchi & Copelli, 2006) and information transmission and storage capacity (Shew et al., 2011). Notably, brain activity near the phase transition is characterized by intermediate synchronization levels but high variability which has been shown both in modelled (Cabral et al., 2011) and empirical data (Fosque et al., 2022; Fuscà et al., 2023). One proposed characteristic of critical dynamics is the existence of metastable states in the brain (Tognoli & Kelso, 2014), which is supported by both experimental and modelling studies (Cabral et al., 2022; Castaldo et al., 2023; Deco, Kringelbach, et al., 2017; Deco & Jirsa, 2012; E. C. A. Hansen et al., 2015). Importantly, metastability has been linked to transient or damped oscillations in that oscillators influence each others’ lifetimes through transient periods of synchronization (Hancock et al., 2022; Torres et al., 2024); thus, metastability arising from delayed interactions between oscillators may be crucial to the integration-segregation balance in the brain (Cabral et al., 2022).

We here establish that those parameter combinations in the whole-brain model which were characterized by intermediate mean values, but high standard deviations of FC strength showed the best fits with FC in source-reconstructed MEG data. In contrast, fits between model and experimental MEG data were less good for “hypersynchronized” modelled data, and even partially negative when controlling for SC as a covariate. These findings were similar between the two coupling modes, PS and AC, albeit with slightly better fits and wider “hypersynchronized” and “high-variability” regions for PS. These results thus provide evidence that both coupling modes of MEG connectivity - phase synchronization and amplitude coupling - emerge near a phase transition in a regime of high variability and metastability.

### Functional connectivity explains MEG beta-band oscillatory network dynamics

The best MEG network - model fits were observed roughly in the range of the high-alpha and beta bands but for low coupling strengths also for low-gamma band up to 40 Hz, i.e., the oscillator’s natural frequency. This was somewhat surprising, since as observed in previous research (Siebenhühner et al., 2024; Zhigalov et al., 2017), the MEG resting-state oscillation connectivity networks peaked in the alpha band for which the model - MEG fit were lower than for beta. This was observed especially when controlling for SC as a covariate, and across a wide range of realistic delays and small-to-medium overall coupling strengths as well as when assessing fits between the model and MEG filtered at different frequencies each. First, this provides evidence that, as proposed by biophysical models (Cannon et al., 2014; Hughes & Crunelli, 2005), oscillatory networks in beta and alpha bands are caused by different neurophysiological mechanisms, of which the beta band networks are more precisely modelled by the Hopf model than alpha or other frequency bands. This is intriguing as the Hopf model and other models are inspired by the biophysical network models, which suggest that synaptic interactions among excitatory pyramidal neurons (PNs) and GABAergic inhibitory interneurons (INs) form the simplest universal microcircuit that can intrinsically generate transiently synchronized oscillations through recurrent and reciprocal interactions (Cannon et al., 2014; Traub et al., 2004). However, the model fits being weak for the alpha-band FC suggest that they originate from a different mechanism that is not modelled by transient oscillations in the Hopf model. Such a mechanism could for example be GAP-junction mediated alpha waves that have been associated with alpha activity in thalamocortical models (Hughes & Crunelli, 2005).

### Structural connectivity influences both model and MEG beta-band connectivity

It has been well established that fMRI-FC is influenced by SC via structure-function coupling (SFC) (Baum et al., 2020; Castaldo et al., 2023; Griffa et al., 2022; Liu et al., 2023; Preti & Van De Ville, 2019; Sadaghiani & Wirsich, 2020). In contrast, thus far, correlation of MEG connectivity with the underlying structure has been studied less, especially for phase synchronization. Crucially, the best model fits of MEG-FC were obtained when modelling was performed with realistic delays informed by SC, and with small coupling strengths which produced transient oscillations with realistic lifetimes that exhibited high variability in functional connectivity. We here found that SFC in MEG data was the strongest for both PS and AC in beta band, and also strong in delta and theta bands for AC, while for PS, it was considerably lower in theta and around zero in delta. Regarding AC, this is in line with earlier findings that also reported peak SFC in beta (Castaldo et al., 2023; Garcés et al., 2016; Liu et al., 2023; Tewarie et al., 2019), and similar results have been obtained for partial coherence (Wodeyar & Srinivasan, 2022). The stronger SFC in lower frequencies for PS than AC reinforces that PS and AC diverge more in these bands, as reported before (Siebenhühner et al., 2024), pointing at different generating mechanisms for low-frequency phase synchronization.

As the Hopf model is informed by both connection strengths and interaction delays from SC, we assessed whether SFC could explain the observed fits between model and MEG FC. Crucially, even after SFC was partialled out, the model prediction remained robust in beta band for PS, being slightly weaker for AC. These data underscore that model dynamics play a pivotal role in explaining experimental MEG networks beyond simple SFC. The differences in MEG-model fits between PS and AC networks demonstrate the partially different mechanistic origin of phase-synchronization and oscillation amplitude coupling (Engel et al., 2013; Siebenhühner et al., 2024; Siems & Siegel, 2020) which may include different influences of neurotransmitter systems (Siebenhühner et al., 2024).

Interestingly, the correlation of model FC with SC was high in both the “hypersynchronized” regions of parameter space, and most of the “high-variability” regions around the former. Moreover, in the “hypersynchronized” regions, the fit between model and MEG vastly decreased when SC was used as a covariate, sometimes even becoming negative, whereas, in the “high-variability” regions, the fit model-MEG remained positive, albeit diminished, with SC as a covariate. This implies that SC generally had a strong influence on modelled FC as in previous studies using the same model (Castaldo et al., 2023), but that only model FC with high variability, i.e., metastable FC, strongly resembled real FC in MEG. Together, these results indicate that neither the high structure-function coupling for the MEG beta band, nor metastability, can alone explain MEG FC, but that both these complex dynamics features are necessary for MEG FC to emerge. This interpretation is also compatible with a novel conceptualization of (brain) criticality where characteristics of critical dynamics are not limited to a narrow point, but stretched into an extended regime - the Griffiths Phase which emerges due to heterogeneity in SC (Moretti & Muñoz, 2013; Ódor & de Simoni, 2021) and determines regional, spectral, and inter-individual variability of MEG PS (Fuscà et al., 2023).

Future studies are needed to discover how laminar asymmetries in feedforward and feedback connections (Castaldo et al., 2023; Deco et al., 2021; Wang, 2020) and E/I homeostasis ((Castaldo et al., 2023)) might improve models and enable better prediction of emergent brain dynamics.

## Methods

### MEG participants

We analyzed here resting-state data from 60 subjects (29 females, mean age 30.6 ± 8.5 years) that had been recruited for various neuroimaging studies at Helsinki University Hospital. All participants reported no recent history of neurological or neuropsychiatric illness and were physiologically eligible to undergo both MEG and MRI measurements. All subjects gave written consent prior to the studies which were approved by the Ethical Committee of the University of Helsinki and performed according to the Declaration of Helsinki.

### MEG recordings and preprocessing

Resting-state MEG (306 channels) was recorded with a Triux system (Elekta-Neuromag) at the BioMag Laboratory, Helsinki University Hospital. Recordings lasted 10 minutes and subjects were instructed to sit still and fixate on a cross in front of them. Temporal signal space separation (tSSS) in the Maxfilter software (Elekta-Neuromag) was used to suppress extracranial noise from MEG sensors and to interpolate bad channels.

We used independent components analysis (ICA) adapted from the MATLAB toolbox Fieldtrip, http://www.fieldtriptoolbox.org/, to extract and identify components that were correlated with ocular artifacts (identified using the EOG signal), heartbeat artifacts (identified using the magnetometer signal as a reference), or muscle artifacts.

### MRI recordings and colocalization

Individual T1-weighted MRI images were recorded from all subjects with a 1.5 T scanner (Siemens, Germany) using a MP-RAGE protocol with a resolution of 1×1×1 mm. MNE software (https://mne.tools/stable/index.html) (Gramfort et al., 2014) was used for the preparation of cortically constrained source models for MEG–MRI colocalization, forward and inverse operators. The source models had dipole orientations fixed to pial-surface normals and a 5-mm inter-dipole separation throughout the cortex, where hemispheres had between 5080 – 7645 active source vertices.

### Source reconstruction and filtering

We then obtained fidelity-weighted collapsing operators (Siebenhühner et al., 2020) that serve to increase reconstruction accuracy when collapsing source time series into the cortical parcels of the Schaefer atlas (Schaefer et al., 2018). After source-reconstruction of MEG data, the resulting parcel time series were filtered into narrowband time series by convolution with a bank of 41 Morlet filters with wavelet width parameter m = 5 and approximately log-linear spacing of center frequencies f_filt_ ranging from 1.1 to 95.6 Hz. Modeled time series were filtered with the same Morlet filters as MEG data.

### Structural connectome

Matrices of structural connectivity were computed from diffusion spectrum and T2-weighted Magnetic Resonance Imaging (MRI) data obtained from 57 participants scanned at the Massachusetts General Hospital centre for the Human Connectome Project (HCP). The connectivity matrix *C* was obtained by counting the number of fibres detected between each pair of N = 100 cortical parcels defined in the Schaefer parcellation scheme. Similarly, the distance matrix *D* was obtained by computing the mean length of all fibres detected 318 between each pair of parcels.

### Model

We used here the whole-brain model of coupled Stuart-Landau oscillators near a Hopf bifurcation with added Gaussian noise (Cabral et al., 2022; Ponce-Alvarez & Deco, 2024). In this model, the interaction between local oscillation dynamics and global coupling (using pairwise between-region coupling strengths *C*_xy_ and delays τ_xy_) interact to produce metastable modes of synchronized transient oscillations near a Hopf bifurcation. Specifically, transient or dampened oscillations emerge in the subcritical Hopf regime (when the bifurcation parameter a is just below 0). Notably, the frequencies of the observed transient oscillations depend on model parameters but tend to be lower than the S-L oscillators’ natural frequency *f*_*nat*_ (Cabral et al., 2022).

Matching our MEG data, we here chose to use the 100 cortical parcels of the Schaefer parcellation as the brain regions in the model, using the C and D connectivity matrices from the HCP to inform the *C*_xy_ and τ_xy_.

The exact temporal evolution of the oscillators is described by the following equation

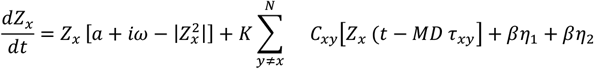

where *Z*_*x*_ is the complex time series of the *x*th oscillator; *a* is the Hopf bifurcation parameter; *ω*=2*π*\**f*_*nat*_ is the angular frequency, with *f*_*nat*_ being the oscillators’ natural frequency; *K* is the global coupling scaling factor, *MD* is the mean conduction delay scaling factor; *C*_xy_ is the connectivity strength between oscillators *x* and *y* taken from the HCP-derived C matrix; *τ*_xy_ is the conduction delay between *x* and *y* taken from the HCP-derived D matrix; *η*_1_ and *η*_2_ are noise terms independently drawn from a Gaussian distribution with mean zero and standard deviation, and the noise scaling factor *β* = 0.001. All modelled time series had a length of 50 seconds at 500 samples per second.

### Assessing oscillations with the phase autocorrelation function

The phase autocorrelation function (pACF) was recently introduced as a way to assess “rhythmicity” of time series (Myrov et al., 2024). We computed, for each parcel and each f_filt_, mean pACF lifetimes, indexing the average lifetime of oscillations in both model and MEG time series.

### Estimation of phase synchronization and amplitude coupling

We estimated pairwise synchronization between all 100 parcels, at all filtering frequencies, using the phase-locking value (PLV) (Tass et al., 1998) for model data and imaginary PLV (iPLV) for MEG data. PLV is sensitive to zero-lag false-positive interactions which arise due to linear mixing in MEG and EEG data (J. M. Palva et al., 2018). We started by computing the complex PLV as

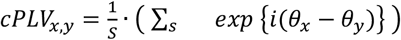

where *θ*_*x*_ and *θ*_*y*_ are the instantaneous phase time series of the complex analytical narrowband time series *Z*_*x*_ and *Z*_*y*_, and *S* is the number of samples s. From this, we derived:

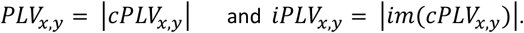

We then estimated, for each frequency and each parcel pair, the group average across all 60 subjects in MEG.

We similarly computed amplitude coupling between all parcels, at all frequencies, using the correlation coefficient (CC) in model data, and the orthogonalized CC (oCC) in MEG data. Similar to iPLV, oCC is insensitive to spurious observations from linear mixing (Brookes et al., 2012; Hipp et al., 2012).

### Fitting modelled data to MEG data

We estimated correlations between group-averaged MEG and model FC (for both PS and AC), using Spearman’s correlation coefficient. This was done both for identical filtering frequencies, and also using individual filtering frequencies for modelled and MEG data.

We used “spin” permutations as an additional test for significance in which parcels, within each hemisphere, are rotated randomly across the spherical surface in the FreeSurfer fsaverage surface. This method preserves in the permuted data spatial autocorrelations which are present in neuroimaging data and have been brought forward as a possible source of false positives (Markello & Misic, 2021). Adapting the procedure described in (J. Y. Hansen et al., 2022), we performed N=1,000 permutations and estimated p-values from the distribution of permuted correlation strengths.

