## Supplemental Figures for "Spectral patterns of MEG oscillatory coupling emerge from meta-stable dynamics with small coupling delays"

### Figure S1

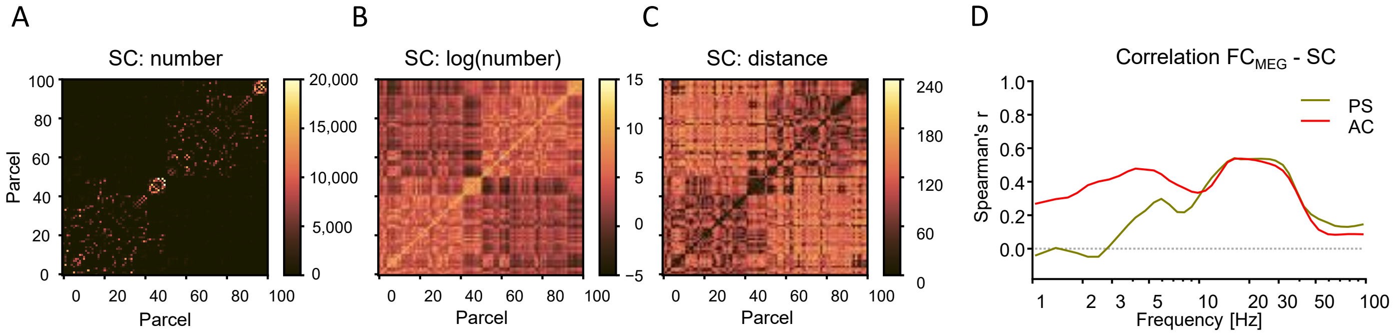


**Figure S1. Structural connectivity.**

**A**. Pairwise strength (number of tracts) of structural connectivity (SC) between 100 parcels (Schaefer atlas) obtained from DTI tractography data (57 healthy subjects from the Human Connectome Project). Self-connectivity was set to zero. **B**. Natural logarithm of SC. **C**. Pairwise distance between the 100 parcels.
**D**. Correlation (Spearman's r) of phase synchrony (PS), and amplitude correlations (AC) in resting-state MEG data (mean over N subjects) with SC.

Figure S2

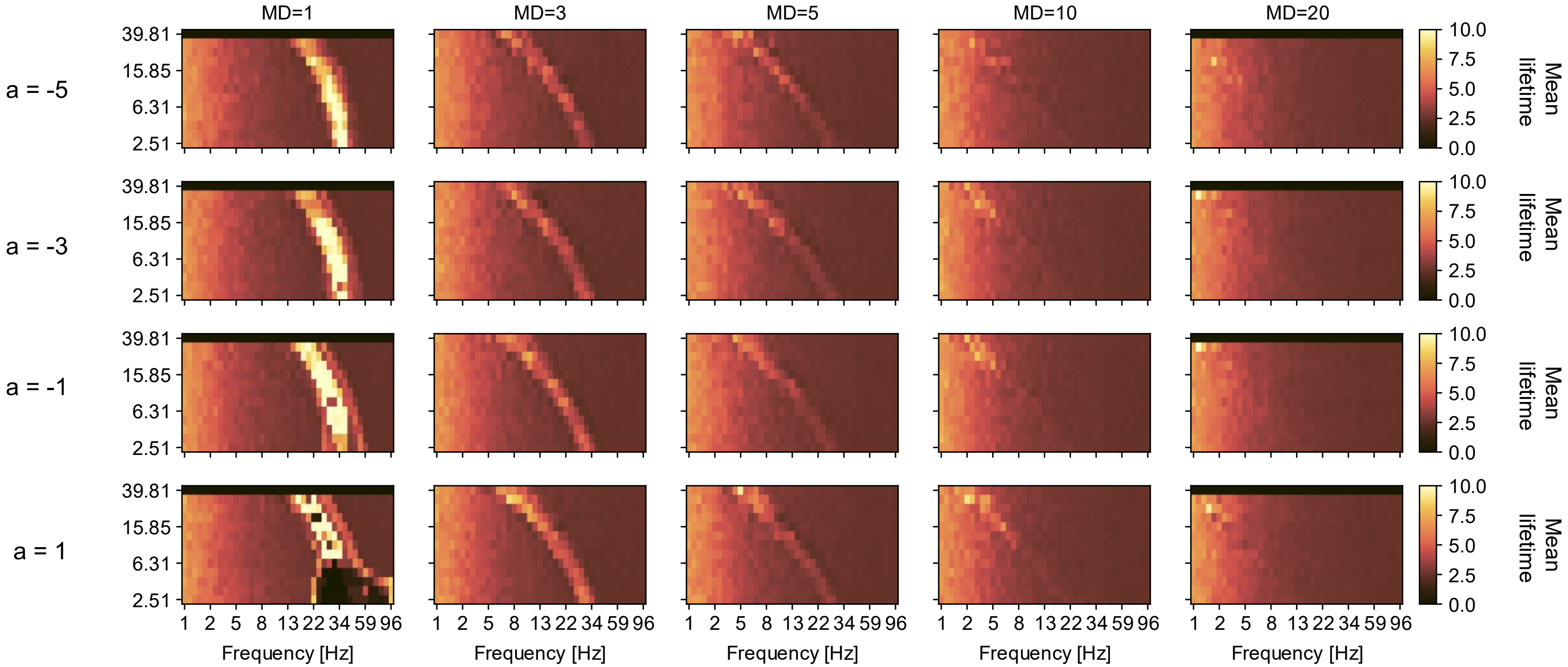


**Figure S2. Phase autocorrelation lifetimes vary with Hopf bifurcation parameter a.**

Oscillation lifetimes, estimated with pACF, as a function of filtering frequency (x-axis), coupling strength K (y-axis), mean delay (MD, columns), and bifurcation parameter a (rows), while natural frequency of the Stuart-Landau oscillators is kept constant at 40 Hz.

Figure S3

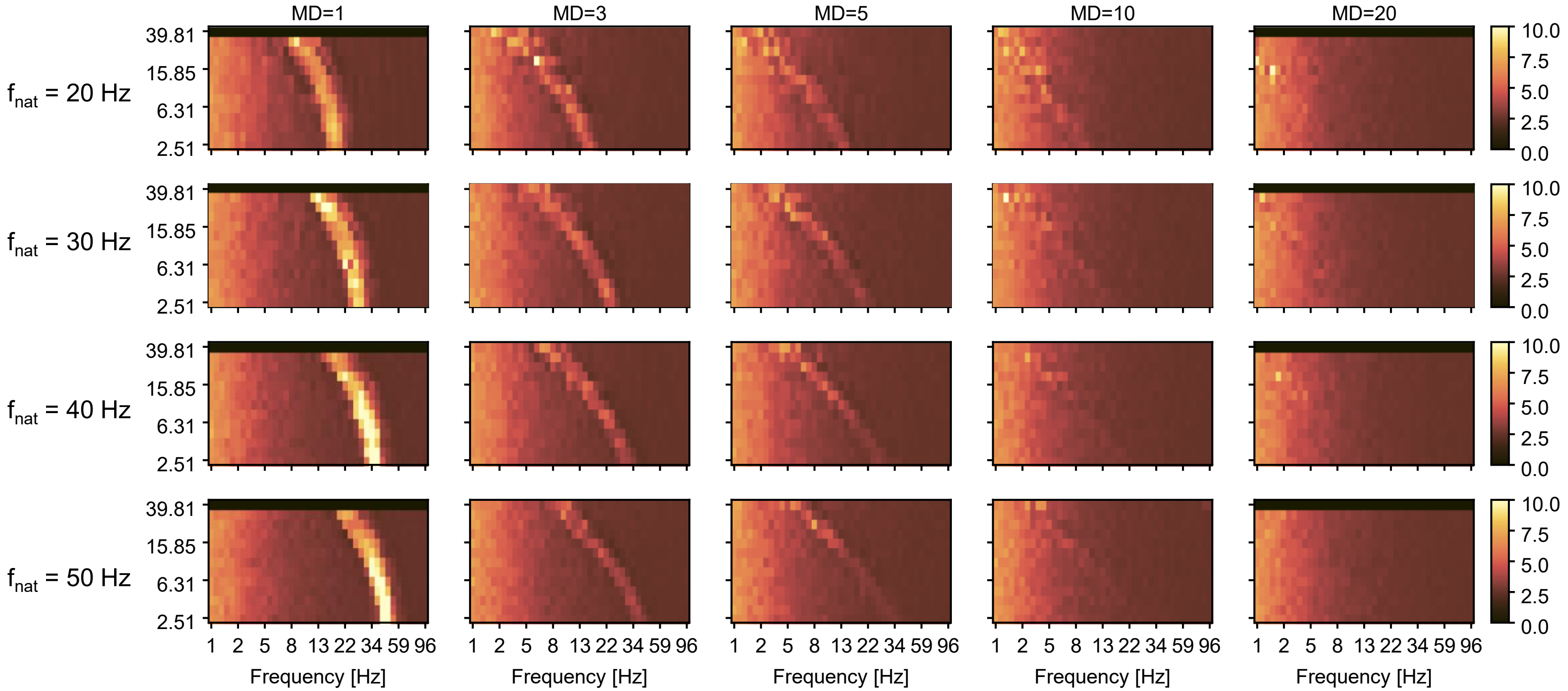


**Figure S3. Phase autocorrelation lifetimes vary with Stuart-Landau oscillator frequency.**

Oscillation lifetimes, estimated with pACF, as a function of filtering frequency (x-axis), coupling strength K (y-axis), mean delay (MD, columns), and natural frequency of the Stuart-Landau oscillators (rows), while Hopf bifurcation parameter a is kept constant at -5.

Figure S4

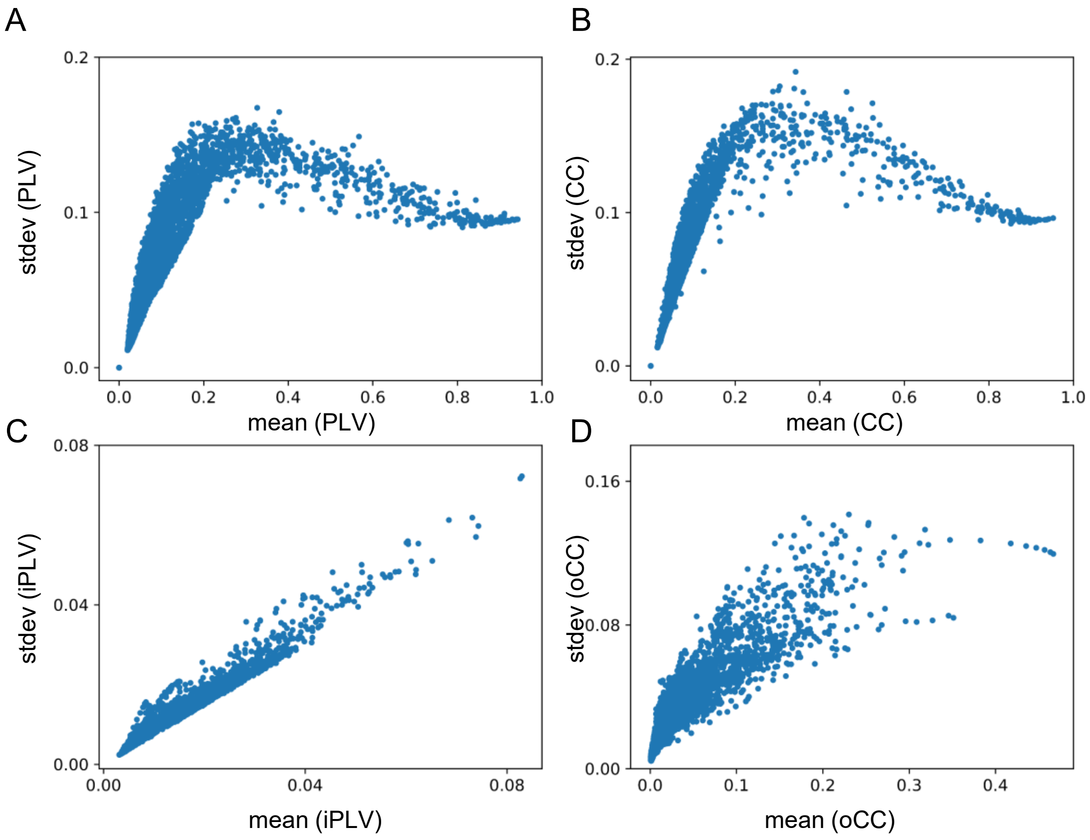


**Figure S4. Functional connectivity in MEG and modelled data.**

**A**. Standard deviation and mean for phase synchrony and **B.** amplitude correlations in modelled data. Each point represents the mean and stdev over 4900 edges between the 100 parcels in the model for a specific combination of K, MD, and f_filt values; with f_nat = 40 Hz and a = -5 in all cases.

**C.** Standard deviation and mean for phase synchrony and **D.** amplitude correlations in MEG data. Each point represents the mean and stdev over 4900 edges between the 100 parcels in the model for a single subject and filtering frequency.

Figure S5

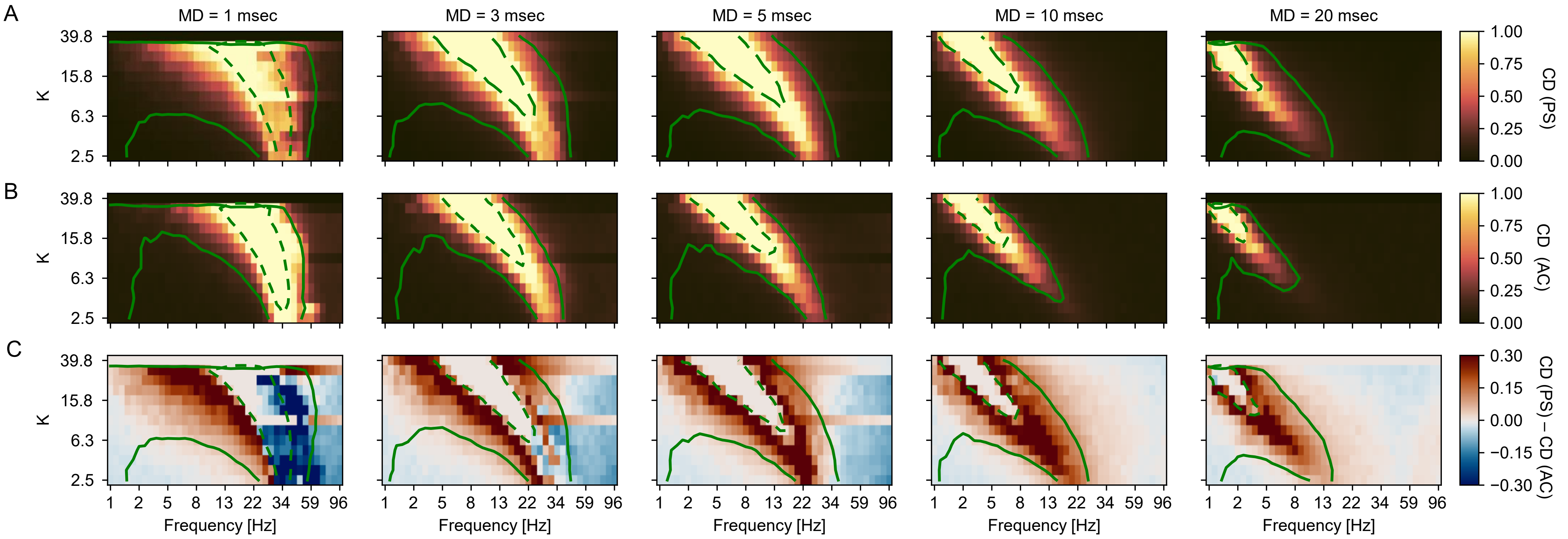


**Figure S4. Connection density of functional connectivity in modelled data.**

**A**. Connection density of phase synchrony (PS) as a function of filtering frequency (x-axis) and coupling strength (y-axis), with a = -5 and $f_{nat}$ = 40 Hz; the color intensity reflects the fraction of edges that are significant ($p < 0.01$) against time-shuffled surrogate data ($N_{perm}$ = 500). Dashed and full contours reflect median values of the mean and standard deviation of PS, respectively, as in Figure 3.

**B**. Same for amplitude coupling (AC).

**C**. Difference in connection density between PS and AC; blue indicates that a higher fraction of edges are significant for PS, red indicates a higher fraction of AC edges.

Figure S6

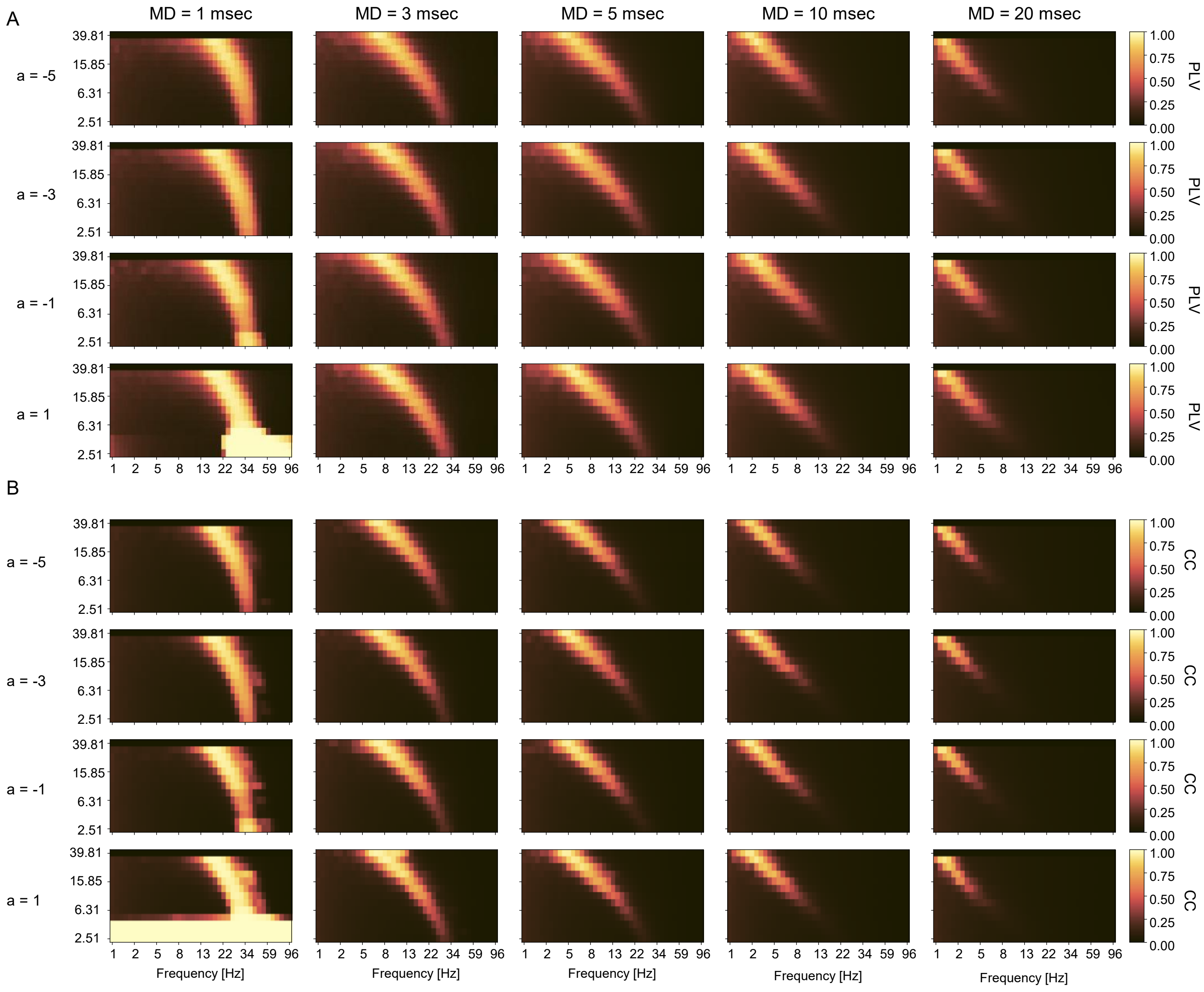


**Figure S6. Mean functional connectivity in modelled data for different values of the bifurcation parameter a.**

**A.** Mean phase synchrony as a function of filtering frequency (x-axis), coupling strength K (y-axis) and mean delay MD (columns), for varying values of the bifurcation parameter a, while $f_{nat}$ = 40 Hz.

**B.** Same as A, for amplitude coupling.

Figure S7

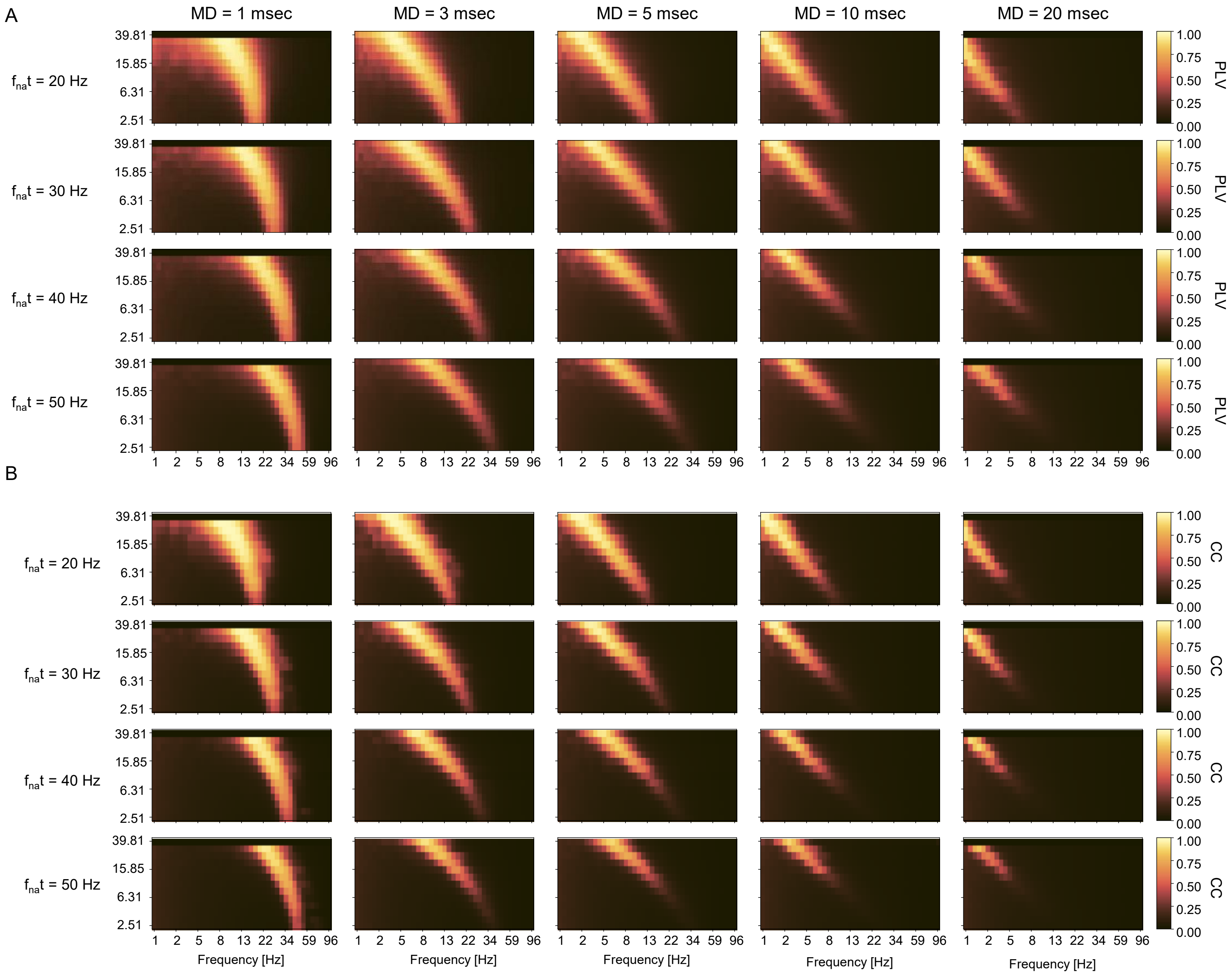


**Figure S7. Mean functional connectivity in modelled data for different values of the oscillators' natural frequency $f_{nat}$.**

**A.** Mean phase synchrony as a function of filtering frequency (x-axis), coupling strength K (y-axis) and mean delay MD (columns), for varying values of the natural oscillator frequency $f_{nat}$ = 40 Hz, while a = -5.

**B.** Same as A, for amplitude coupling.

Figure S8

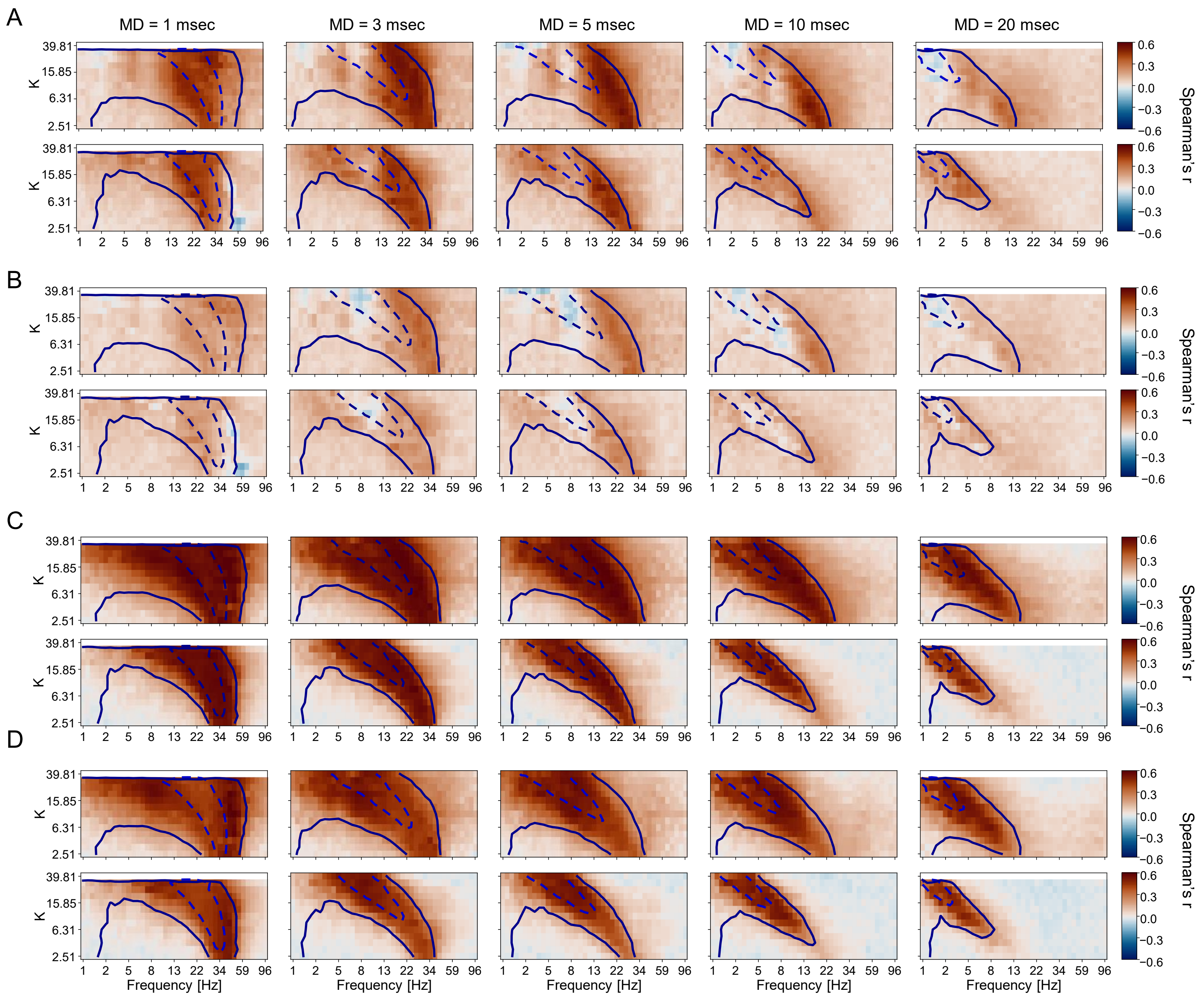


**Figure S8. Correlations between modelled and MEG data and between modelled data and SC.**

**A.** Spearman's correlation (over all edges) of modelled FC with MEG FC as a function of filtering frequency (x-axis), coupling strength K (y-axis) and mean delay MD (columns), for a = -5 and $f_{nat}$ = 40 Hz. Dashed and full contours reflect median values of the mean and standard deviation of PS, respectively, as in Figure 3.

**B**. Same, using SC as a covariate.

**C.** Spearman's correlation (over all edges) of modelled FC with SC as a function of filtering frequency (x-axis), coupling strength K (y-axis) and mean delay MD (columns), for a = -5 and $f_{nat}$ = 40 Hz.

**D**. Same, using MEG FC (at same filtering frequency) as a covariate.

Figure S9

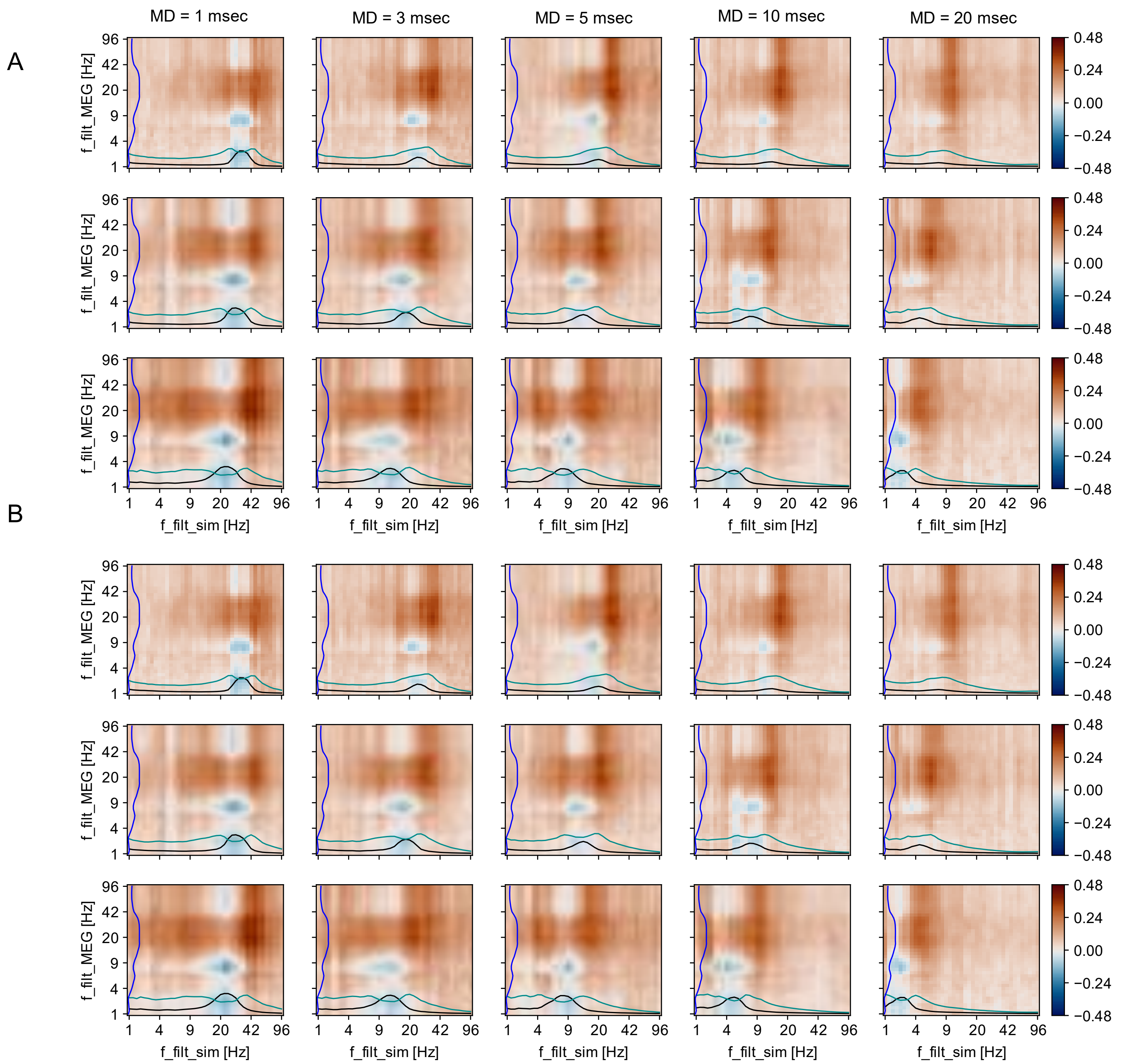


**Figure S9. Correlation between model and MEG across frequencies, with structural connectivity as covariate.**

**A.** Correlations (Spearman's r, across all edges) between model PS and group-mean MEG PS when both are filtered at different Morlet frequencies (x-axis: $f_{filt}$_model and y-axis: $f_{filt}$_MEG) for different values of K (rows) and MD (columns), with structural connectivity as a covariate. Black and cyan lines show mean and standard deviation of PS in the model as a function of frequency (a.u.).

**B**. Same for AC.
